# Novel regression methods for metacognition

**DOI:** 10.1101/423947

**Authors:** Simon Bang Kristensen, Kristian Sandberg, Bo Martin Bibby

## Abstract

Metacognition is an important component in basic science and clinical psychology, often studied through complex, cognitive experiments. While Signal Detection Theory (SDT) provides a popular and pervasive framework for modelling responses from such experiments, a shortfall remains that it cannot in a straightforward manner account for the often complex designs. Additionally, SDT does not provide direct estimates of metacognitive ability. This latter shortcoming has recently been sought remedied by introduction of a measure for metacognitive sensitivity dubbed meta-*d*’. The need for a flexible regression model framework remains, however, which should also incorporate the new sensitivity measure. In the present paper, we argue that a straightforward extension of SDT is obtained by identifying the model with the proportional odds model, a widely implemented, ordinal regression technique. We go on to develop a formal statistical framework for metacognitive sensitivity by defining a model that combines standard SDT with meta-*d*’ in a latent variable model. We show how this agrees with the literature on meta-*d*’ and constitutes a practical framework for extending the model. We supply several teoretical considerations on the model, including closed-form approximate estimates of meta-*d*’ and optimal weighing of response-specific meta-sensitivities. We discuss regression analysis as an application of the obtained model and illustrate our points through simulations. Lastly, we present `R`-software that implements the model. Our methods and their implementation extend the computational possibilities of SDT and meta-*d* and are useful for theoretical and practical researchers of metacognition.

## Introduction

Insight into one’s actions and abilities is often termed metacognition. Metacognition is a recurrent concept in many psychological fields of study (Metcalfe & Shimamura, 1996), and metacognitive disability is a characteristic of clinical diagnoses such as schizophrenia, Alzheimer’s and certain injuries of the brain (David, Bedford, Wiffen, & Gilleen, 2012). In cognitive science, these processes and disorders are investigated in a neurological context through cognitive and behavioural experiments, usually involving a complex setup with repeated measurements of test subjects over various stimulus intensities and possibly other changes of experimental settings such as blocked trial runs. The basic structure of most such experiments can be summarised as follows: In a single trial replication, a subject must first complete a task (e.g. identify which of two images were presented) and thereafter report a quantification of her/his confidence in, awareness of, or feeling of control over the performed task. The first assignment is usually dubbed the type 1 task while the reflection over the type 1 task is called the type 2 task. Various quantifications in the type 2 task have been proposed, such as verbal and numeric rating scales of confidence (Dienes, 2007; Dienes, Altmann, Kwan, & Goode, 1995), awareness (Ramsøy & Overgaard, 2004; Sandberg, Timmermans, Overgaard, & Cleeremans, 2010), or as a probability assessment (Dienes, 2007; Fleming & Daw, 2017), and more indirect methods have been proposed, for example allowing the subject to wager money on the correctness of the type 1 assertion (Kiani & Shadlen, 2009; Persaud, McLeod, & Cowey, 2007).

A common assumption when modelling the responses of consciousness experiments is that the subject’s decision is based on thresholds in an underlying evidence distribution so that the response changes when the signal crosses a threshold, a model referred to in psychology as Signal Detection Theory (SDT) (Green & Swets, 1966). While the signal detection model remains ubiquitous in modelling cognitive experiments, the question of how to use it in quantifying metacognitive capabilities has been revisited and revised multiple times, see Fleming and Lau, 2014 for a review. Additionally, later years have seen various expansions and alternatives to the SDT, some examples of which can be found in Maniscalco and Lau, 2016. Such models may be termed ‘formal’ as they are meant to represent the underlying, e.g. neural, process that the researcher believes to explain the response in a single trial, while they remain abstracted from design structure, data acrual and methods of estimation.

The formal model will specify a relationship between the processes controlling performance and insight leading to some index of metacognitive ability. A recently proposed measure in the SDT setting is meta-*d* (“meta d prime”, Maniscalco & Lau, 2012; Rounis, Maniscalco, Rothwell, Passingham, & Lau, 2010), meant to quantify a subject’s sentivity to the difference between a correct and incorrect response. The introduction of meta-*d* was in itself an attempt to remedy a shortfall of SDT in that the SDT model does not give a direct measure of metacognitive abilities. However, the literature tends to focus on defining meta-*d* from the observed response frequencies and not based on an underlying model.

An inadequacy shared by SDT and most other formal models is that they are not sufficiently elaborate to directly include the complex designs of most cognitive experiments, particularly it is not clear how to account for multiple stimulus intensities or repeated measurements, and it is not obvious how to include experimental factors and subject characteristics as covariates to adjust comparisons or perform predictions. In neuropsychological experiments, we may require for example to regress cognitive abilities on a structural brain measure while adjusting for age and possibly modulate the effect by brain region. These shortfalls are often addressed by applying the SDT multiple times under various dichotomisations and stratifications with substantial loss of efficiency as a consequence. Moreover, failure to account for sources of variation aside from that of the neurological process can lead to biased estimates and erronous inference. Morey, Pratte, and Rouder, 2008, for example, showed how three classic results in recognition memory can be produced as artifacts of ignoring extra-variational factors such as task difficulty.

In the present paper, we analyse models for metacognition from a theoretical perspective focusing on SDT and meta-*d* and compare them to established statistical models. We define a model integrating SDT and the meta-*d* measure and show how this leads to expansions to regression models meant to address the above mentioned shortcomings. These approaches are exemplified through simulations to illustrate the importance of accounting for variational sources in an efficient manner.

While our approach is teoretical, we aim to primarily address a practical, fundamental problem of a researcher planning an analysis of a consciousness experiment: How to convert a formal model to an appropriate statistical analysis. A similar idea underlies Wright, Horry, and Skagerberg, 2009 where the authors consider a simpler experiment with only a type 1 task and propose mixed model techniques to enable statistical analysis in type 1 SDT. First, the coupling of a statistical and formal model has the practical advantage that it allows for computation of estimates and standard errors from sound statistical principles that can, ideally, be performed by standard software. Second, and more theoretically, an understanding of the corresponding statistical model may help to make explicit the underlying assumptions of the formal model and standard techniques for model validation can be applied. Furthermore, the statistical model may offer more extensions which can be converted to insights into the subject matter.

The paper is structured as follows. We begin by recounting the basis of Signal Detection Theory with a vu to some earlier versions and expansions of the basic model. We then establish the relationship between generalised linear models (GLM’s) for binary data and latent variable models, and we recall two statistical models, the proportional odds (PO) model and a natural extension, the partial proportional odds (PPO) model. We then derive the simple, but often overlooked fact that the SDT model for metacognition is equivalent to the PO model (DeCarlo, 1998), a well-known regression tehnique for ordinal data.

Next, we introduce meta-*d* through a statistical model using latent variables and argue that this is in concordance with the definition in for example Barrett, Dienes, and Seth, 2013. We compare the obtained model, dubbed the meta-SDT, to the PPO to show that while the two are conceptually very similar, they differ in an important way that bears consequence to the interpretation of the model parameters. In addition, we discuss model validation and a technique for obtaining approximate, closed-form estimates (i.e. an explicit expression) of the meta parameters. The obtained methods are illustrated and tested through simulations. Finally, we discuss numerical issues connected to the application of regression models for meta-SDT and introduce the `R`-package `metaSDTreg`.

The novel contributions made by this article include a statistical description of meta-*d*’ that is formal in the sense that it connects a specified model (specified through latent variables) directly to meta-*d* in the model’s parameters. This constitutes a flexible framework that is well suited for expansions of the model. We also supply an optimal weight for combining response-specific meta sensitivities and formulas for approximate closed-form estimates of meta-*d*, something that is not to our knowledge presented elsewhere. Additionally, we present software to fit the meta-SDT model coded as a package in the open-source statistical software `R` (R Core Team, 2016).

We end this introduction by introducing some ubiquitous notation. We imagine in the following a cognitive experiment such as that which is presented in Sandberg et al., 2010, but with a two-choice type 1 task. Thus, a subject must for the type 1 task for example report whether a shape was present on an image shown. We denote by *S* ∈ {0,1} the true status with *S* =1 meaning “present” (or “signal”) and *S* = 0 meaning “absent” (or “noise”). The stimulus intensity is varied for example by presenting the image for *t* milliseconds. In the type 2 task, the subject must quantify perceived awareness of the decision using an ordinal scale {1, 2,…, *L*}. We will use the subscripts 𝓝 and 𝓢 to denote quantities in the type 2 model which pertain to a “noise” or “signal” report in the type 1 task, respectively. Subjects are indexed by *i* = 1, …, *N* and replications are indexed by *j* = 1, …, *M*. To minimise unwieldy notation we will often omit these subscripts and consider a single replication on a single subject.

Throughout, we will by Φ and *ϕ* respectively denote the cumulative distribution and density function of the standard normal distribution. We let ℙ (·) be a probability measure, and let 𝔼 [o], Var (·), and ℂov (·, ·) be the associated expectation, variance, and covariance operators, respectively. We further denote by 𝟙_{·}_ the indicator function.

## Cognitive responses: Signal detection theory

For notational parsimony, the SDT model that we introduce here is the simplest version meant to model experiments with a dichotomous type 2 response. This can be naturally extended to ordinal type 2 assertions by broadening the threshold structure. We end this section by discussing earlier versions as well as possible expansions of the signal distributions.

Signal Detection Theory assumes that responses occur as decisions based on a latent evidence *X*. We will adhere to the classical conventions of SDT and assume that *X* follows a standard normal distribution and that a signal shifts the location of the noise distribution by *d* (“d-prime”), but naturally various alternatives and extensions exist (e.g., DeCarlo, 2010).

The SDT model is shown in Figure 1. Responses are governed by a threshold model on *X* specifying that a subject responds “signal” if *X* > *θ* and “noise” if *X* ≤ *θ*. The type 2 response is then “confident” if *X* > *τ*_𝒮_ (meaning confident in signal) or *X* ≤ *τ*_𝒩_ (confident in noise) and otherwise the assertion is “unsure”. One assumes that *τ*_𝒩_ < *θ* < *τ*_𝒮_ under the rationale that for a given response one requires stronger evidence in the given direction to move from “unsure” to “confident”. We refer to *θ* as the type 1 criterion, to *τ*_𝒩_ and *τ*_𝒮_ as the type 2 criteria, and to *d* as the signal sensitivity.

**Figure 1.**
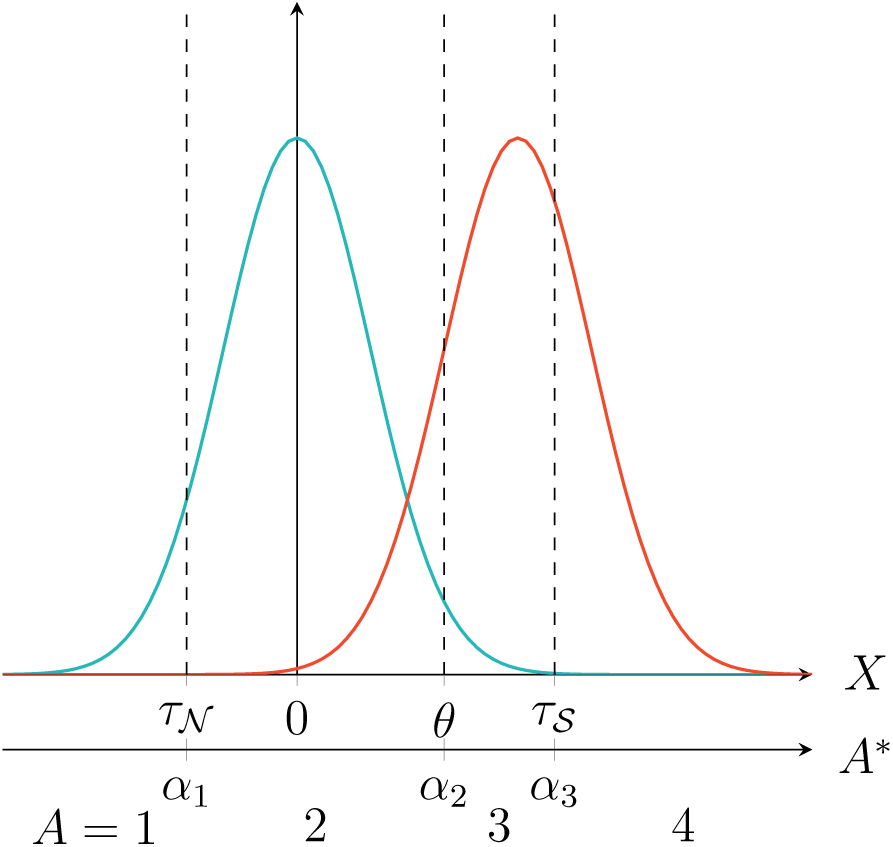
The SDT model with ordinal responses *A*. The black / blue curve is the evidence density under noise (*S* = 0) while gray / red is the evidence distribution under signal (*S* =1). *X* is the underlying evidence governing the responses and *A*\* governs the concatenated, ordinal answers *A*.

Estimation in SDT is usually based on simple relations between probabilities and parameters. Thus one would estimate

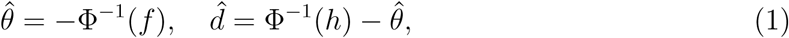

where *f* is observed *F*, the type 1 false alarm rate (the probability of incorrectly reporting a signal), and *h* is the observed *H*, the type 1 hit rate (correctly reporting a signal).

We refer to (1) as moment estimators. In the single-task SDT (i.e. where there is only a type 1 task) the moment estimators coincide with the maximum likelihood estimates when responses are binomial (Appendix A), but this is generally not true when considering the SDT model in Figure 1 and better estimates (in terms of efficiency) are obtained from the proportional odds model (see below), which is available in any standard statistical software package.

### Predecessors and expansions

An important influence for SDT in metacognition is the paper by Galvin, Podd, Drga, and Whitmore, 2003. Galvin and colleagues formulated a response model, where a subject must decide on the type 1 task based on an evidence distribution and then perform the type 2 task using a transformation of the same evidence by some function *ω*. The idea is to enable overall modeling of type 1 and type 2 response ROC curves, an idea which may be said to go back to an editorial letter by Clarke, Birdsall, and Tanner, 1959.

This is contrasted by the SDT model above which more naturally facilitates models for the type 2 task conditioned on type 1 (e.g. confidence given a “signal” response). We will revisit these conditionals below when we discuss how to quantify metacognitive sensitivity. Note also that the approach of Galvin et al. is very similar to the present SDT model taking *ω*(·) = |*θ* – ·|, see Maniscalco and Lau, 2014 for a more in depth discussion. As we will exemplify with the Independence model below, the two approaches may be said to differ in their threshold models.

An obvious alternative to the approach of Galvin et al. and an extension of SDT would be instead to consider a two-dimensional evidence distribution for type 1 and 2 responses. In consciousness and behavioural research, one such model proposed for memory research is the Stochastic Detection and Retrieval model of Jang, Wallsten, and Huber, 2012. A bivariate model for source monitoring is found in DeCarlo, 2003 while another, Bayesian, approach is explored by Fleming and Daw, 2017.

A special case, as we argue below, of the two-dimensional signal distribution is the so-called hierarchical model (Maniscalco & Lau, 2016; Rausch & Zehetleitner, 2017), wherein one assumes that the type 1 task follows a single-task SDT and that the type 2 decision is made in an SDT based on evidence *Y*, where

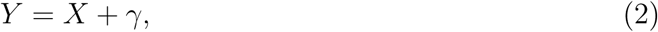

for *γ* ∼ *N*(0, *ζ*^2^) independent of *X*. The reasoning is that *ζ*^2^ measures the noise or loss of evidence in making the type 2 assertion compared to the type 1 task, where *ζ*^2^ = 0 recovers the SDT.

The relationship between the distributional formal models are shown in Figure 2. This is, however, only the signal distributions and one must also specify the threshold model. The best example of the importance of thresholds is the independence model (Rausch & Zehetleitner, 2017). If we in the independence model were to specify the thresholds in the spirit of Galvin et al., a subject would report a signal if *X* > *θ* and confidence if *Y* > *τ* yielding a rather odd model in which the type 1 and type 2 tasks are independent. However using the ordered rationale of SDT (see Rausch and Zehetleitner, 2017 for details),

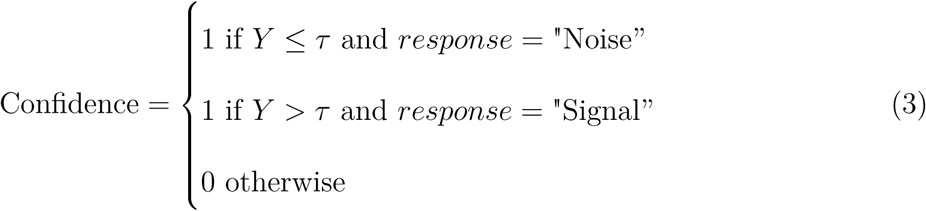

and thus we obtain a model where the probability of confidence (for *Y* symmetrically distributed) given *X* is ℙ (*Y* ≤ (1 – 2𝟙_{*x*>*θ*}_)*τ*), which is not independent of *X*.

**Figure 2.**
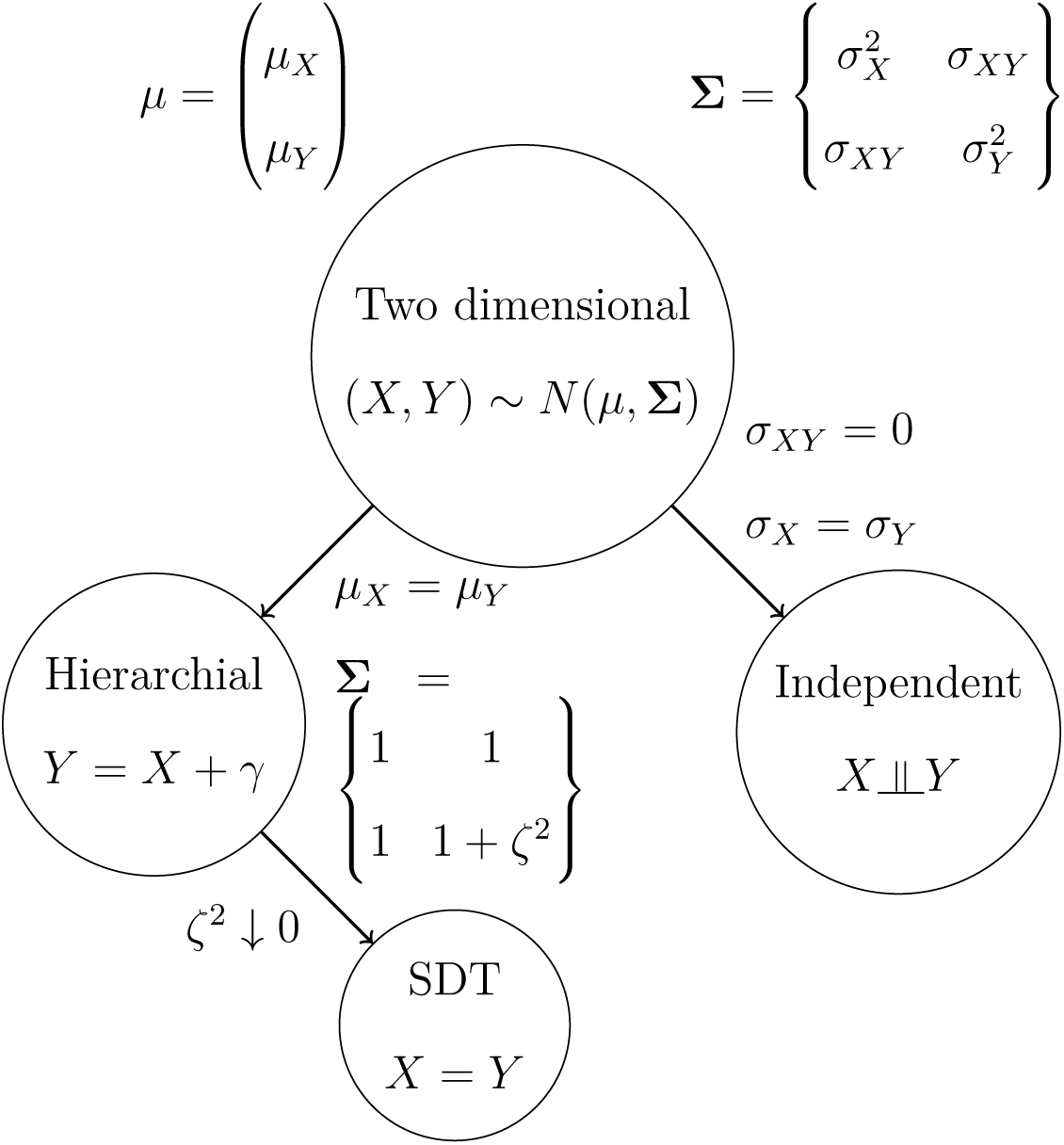
Interrelationship of signal distributions in some common signal models. In the two-dimensional model, the simultaneous distribution of *X* and *Y* is assumed to follow a bivariate normal distribution (note that the variance structure is too general to be identifiable when used to generate latent variables in a threshold model). The other models are obtained as special cases by applying standard results for the multivariate normal distribution. For example, letting the components be independent and equivariable we would obtain the Independent model. Alternatively, taking the mean parameters to be the same and the variance structure additive yields the hierarchical model, of which SDT is the special case for *X* = *Y* occurring as *ζ* goes to zero.

## A brief statistical precursor

In the present section, we recount some of the theory concerning generalised linear models (GLM’s) and introduce two common regression models for ordinal data.

A GLM can be described as a regression for the mean of a response belonging to an exponential family distribution, where it is assumed that the explanatory variables enter additively after transformation using a so called link function (McCullagh & Nelder, 1989). For example, if the response is Gaussian and the link is taken to be the identity function, the GLM is linear regression while a binary response with a logit link would result in logistic regression.

We will focus on binary outcomes with a particular class of link functions. Suppose to this end that *R* ∈ {0,1} is a binary response and that *F* is a distribution function that is differentiable, strictly increasing on the real numbers, and is such that *F* – 1/2 is an odd function. Given covariates *V* we might then consider the GLM

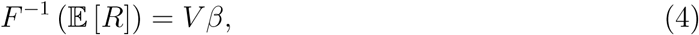

where *β* is a parameter vector. An equivalent formulation of (4) is through a latent variable *R*\* where we associate the latent to the observed by {*R* = 1} = {*R*\* > 0} and specify the latent variable through a regression equation,

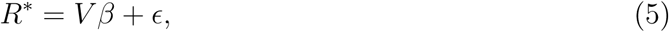

where *ϵ* ∼ *F*. Note that the assumptions on *F* ensure that *ϵ* is absolutely continuous, supported on the real numbers and symmetrically distributed. We will refer to (5) as the latent variable representation of the GLM. Also note that the representation consists of two components, the threshold model linking the latent variable to the observed and a distribution model for the latent variable. The two formulations are indeed equivalent since

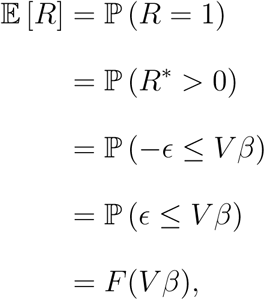

which is equivalent to (4).

The majority of statistical models dealing with ordinal data from an ordered scale {1, …, *K*} can be characterised as a series of binary models that arise from considering cumulative, possibly conditional, probabilities on the scale. The parameters are constricted across models to increase efficiency and parsimony, see Ananth and Kleinbaum, 1997 for a review. Here we will consider the pervasive proportional odds model.

### Proportional odds (PO)

Suppose that we observe an answer *A* ∈ {1, …, *K*} from an ordinal scale. The *proportional odds model* with a probit link is defined as

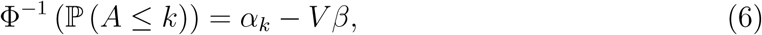

for *k* = 1, …, *K* – 1 and where *α*_1_ < … < *α*_*K*–1_. The sign for *β* is chosen so that the interpretation coincides with that of a standard probit regression when *K* = 2.

The name of the model can be motivated from the following property when using the logit link Λ(*x*) = 1/(1 + exp(–*x*)). If we consider the odds of observing an outcome no greater than *k*_1_, this equals exp(*α*_*k*_l__) exp(–*βV*) and comparing this to the odds of observing an outcome no greater than *k*_2_, the ratio of the two odds is the same for any choice of covariates, the ratio being exp(*α*_*k*_l__ – *α*_*k*_2__). Moreover, say we wish to calculate the change in odds for the event {*A* ≤ *k*} as we change the explanatory variables from *V*_1_ to *V*_2_. This is done using

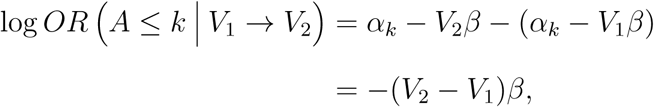

so that the odds ratio is calculated as exp(–(*V*_2_ – *V*_l_)*β*). The assumption of proportional odds expresses that the OR is independent of *k*, i.e. there is no interaction between covariates and scale level.

The proportional odds assumption can be checked by examining so called proportional odds plots, that can be constructed by calculating observed frequencies, transforming to the link scale and plotting these against covariate levels. Deviations from parallel lines will be indicative of violations of the assumption. Other approaches to model validation include investigating partial residuals (Harrell, 2015).

From our considerations on latent variables above, it follows that a latent variable representation of the PO model in (6) is given by {*A* ≤ *k*} = {*A*\* ≤ *α_k_*}, where

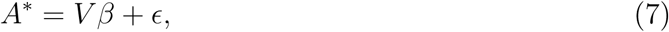

for *ϵ* ∼ *N*(0,1). Note that the proportional odds assumption in the latent variable context expresses that *A*\* does not depend on *k*.

#### The SDT as a PO model

Consider again the SDT model as outlined above. It is clear that the evidence distribution can be represented as *dS* + *X*, which is the same distribution as the latent variable in the PO model (7), since *ϵ* ∼ *X*. Combining this with the fact that the threshold model of the PO model coincides with that in SDT (see Figure 1), it follows directly that SDT is equivalent to the PO model with *K* = 4 and with a single covariate *V* = *S* (in the sense of identical likelihoods). The associated ordinal answers *A* with the latent variable *A*\* are also shown in Figure 1 and the threshold values of the PO model can be associated with those in SDT by

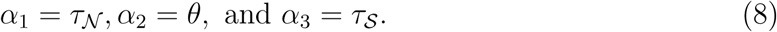

Additionally, the signal sensitivity *d* corresponds to the regression parameter *β* from the PO model.

### Partial proportional odds (PPO)

When the proportional odds assumption seems unrealistic or, for other reasons, a more involved parameter structure is deemed appropriate, a natural extension of the PO model can be defined by relaxing the proportionality assumption for a subset of the covariates *V*_0_,

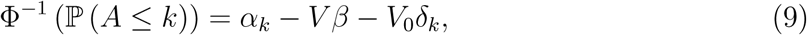

where *δ_k_* measures the deviation from proportionality. The latent variables associated with the PPO are

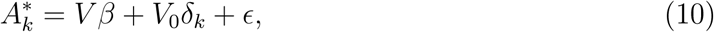

with 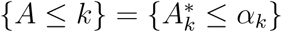 for *ϵ* ∼ *N*(0,1). Naturally, *δ_k_* = 0 for every *k* recovers the PO model.

As an example, we return to our consciousness experiment from the introduction. We define the ordinal variable *A* as a reordering of the responses, see Figure 1. We might wish to let the effect of *S* vary depending on the task type, i.e.

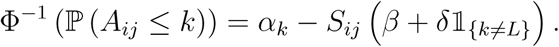

This is the PPO model (9) with *V_ij_* = *V*_0,*ij*_ = *S_ij_* and *δ_k_* = *δ*𝟙_{*k*≠*L*}_. The response to the type 1 task is the event {*A* ≤ *L*} (“noise”) or its complement (“signal”), and this is performed with a discrimination between signal and noise being *β*. When deciding which awareness rating to assign to the type 1 task, this is done with a signal discrimination *β* + *δ*. We argue below that *β* will correspond to the signal sensitivity *d* known from SDT, while *β* + *δ* is similar to, but not the same as, meta-*d*. If we wished to account for between subject variation, we could modify our model as

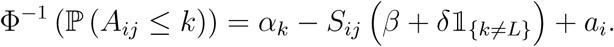

Note that the subject intercepts *a_i_* enter as a proportional effect, i.e. it is assumed that a given subject has a response bias that she/he applies equally to both the type 1 and type 2 assertion. Another specification could be to assume that the subject effects were realisations of some population distribution, in which case one would obtain a mixed model. While it is possible to let the subject effects depend also on *k*, some care is needed as the subject-specific intercepts should still be ordered and reparametrisation is generally needed to ensure ordinality of the scale (Tutz & Hennevogl, 1996). We will term the present model, with no random effects, a fixed effect model.

## The meta-SDT model

Below we introduce the main contribution of the present paper, when we define a model for all responses from a meta-cognitive task which includes in its parameters the response-specific meta-*d*’s which are concordant with the definition in e.g. Barrett et al., 2013. The model relies on a latent variable formulation similar to that in Signal Detection Theory, but also employing the truncated normal distribution 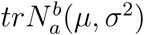 (Appendix B).

We discuss first an intuitive definition of meta-*d* before moving on to the meta-SDT model. This is presented first in the simple case with a two-level type 2 rating and a binary signal intensity. In this case, we study the likelihood to compare to the standard SDT model and to the PPO model. Next, we generalise this model to accommodate an ordinal type 2 task and a general regression model in order to account for more involved designs. We end this section by making a few separate, discussive points concerning the obtained model, including a comparison to the PPO model, model validation and how to obtain a joint meta-*d* measure from the response-specific meta-*d*’s. We additionally study how to obtain approximate, closed-form estimates of meta-sensitivity.

### >Meta-*d*

In the same manner as *d* measures the sensitivity to signal in the standard SDT model, meta-*d* is meant to quantify meta-sensitivity, i.e. the individual’s ability to separate correct from incorrect decisions. Meta-*d* was introduced in Rounis et al., 2010 and more thoroughly discussed by Maniscalco and Lau, 2012, while a more mathematical approach is found in Barrett et al., 2013 and Maniscalco and Lau, 2014.

Figure 3 shows the distribution of awareness ratings from an experiment for correct and incorrect answers to a shape recognition task. Note that this is something of a muddling of data as the responses are measured at various stimulus intensities and for different subjects. The overall picture, however, shows a pronounced tendency to score an incorrect answer low on the perception awareness scale while a high score is often given to correct assertions and we might take this as an indication of high meta-sensitivity.

**Figure 3.**
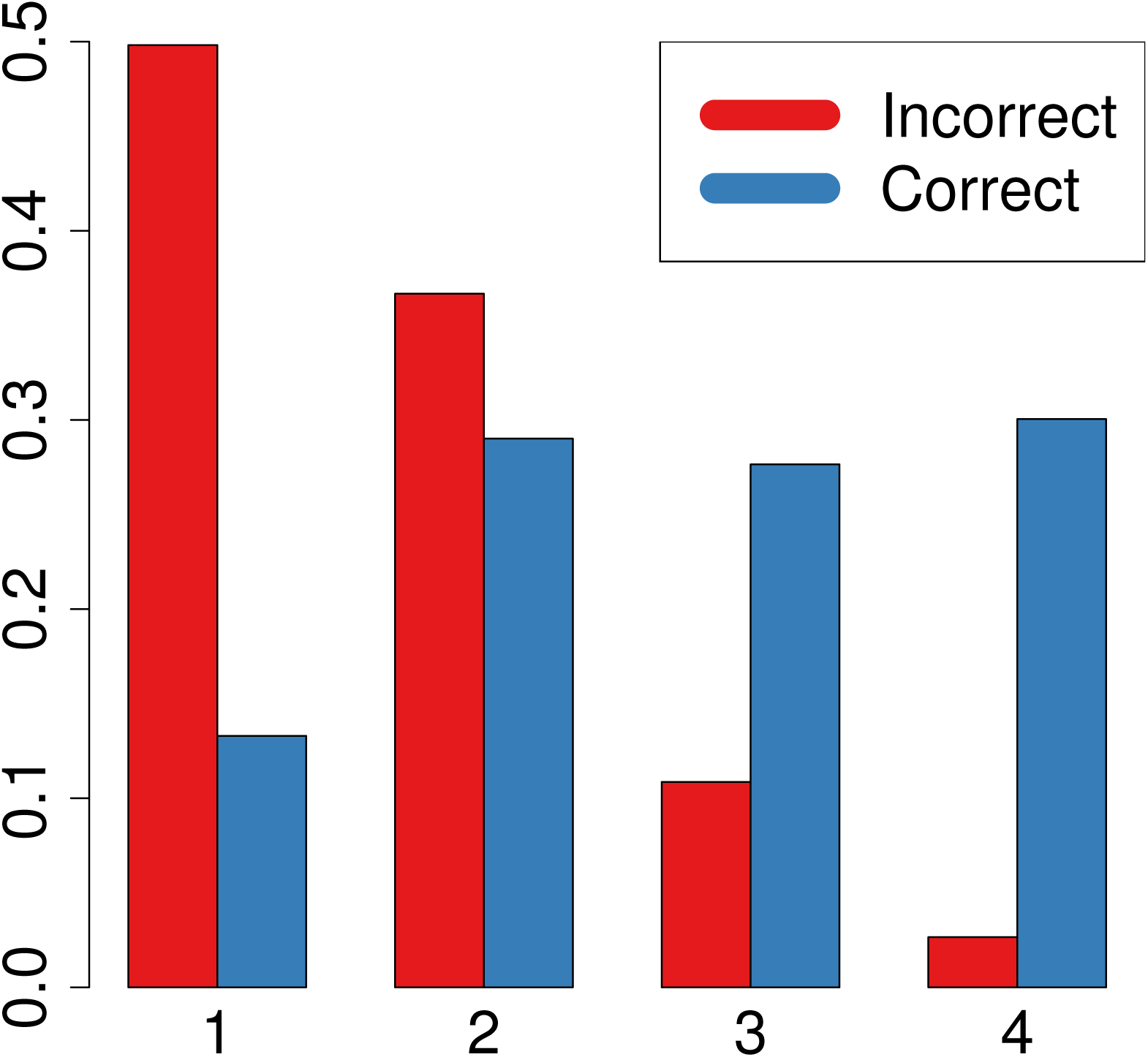
Distribution of Perception Awareness Scale ratings for incorrect and correct assertions.

Initially, we may wish to separate correct and incorrect assertions conditional on whether the type 1 report was “noise” or “signal”, as a difference between response types may confound the aggregate measure. This leads to the concept of response-specific meta-*d*. When defining the response-specific meta-*d*’s in the SDT model we use two basic properties of SDT: That conditional on the type 1 response, the noise and signal distributions represent correct/incorrect response distribitions and that the SDT model implies the response-specific evidence distributions and vice versa (barring some assumptions). This idea is illustrated in Figure 4 for the signal-specific response.

**Figure 4.**
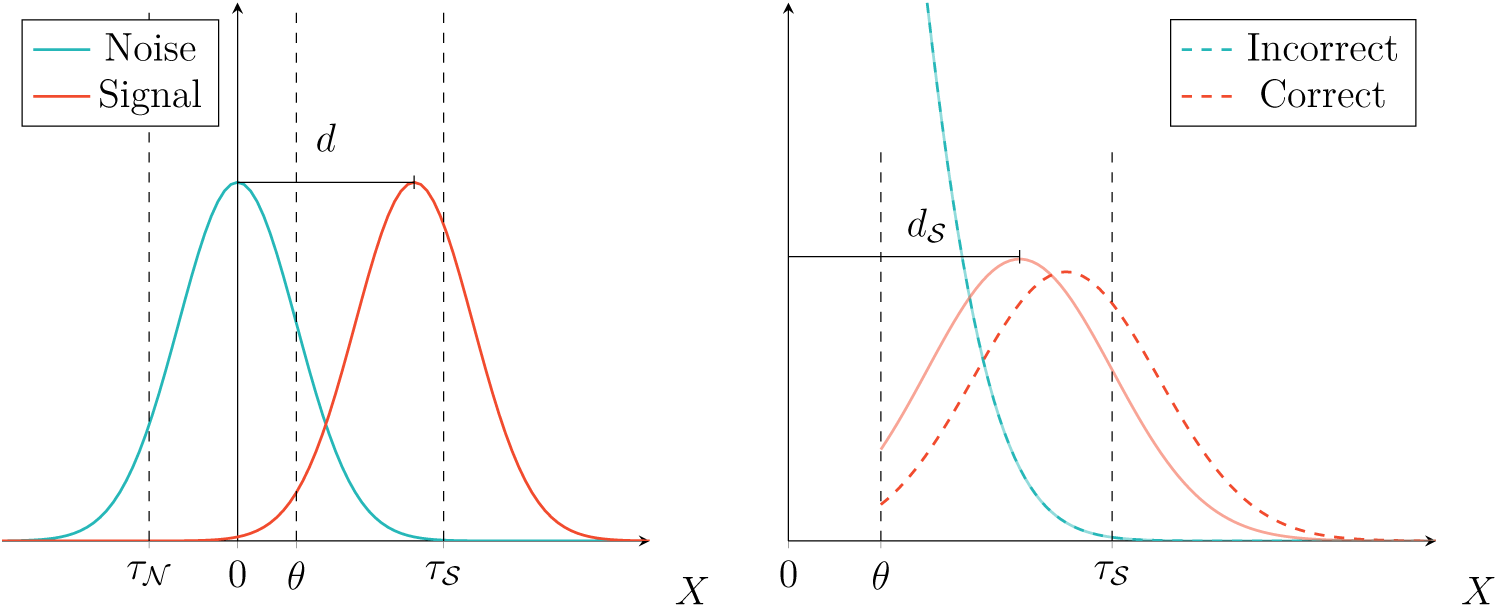
Intuitive definition of meta-*d*: The left plot depicts the SDT model. Conditioning on a “signal” response (*X* > *θ*), we obtain the right hand plot where the dashed lines are those expected if the type 2 decision is based on the SDT. If we conversely start from the signal-specific type 2 data in the right-hand plot we may ask which SDT model has generated these responses (the fully drawn lines). The signal-specific meta-*d*, *d*_𝒮_, is then the sensitivity in this SDT model. Note that the fully drawn lines in the right-hand plot is the density of *Z*_𝒮_ in (12).

### Meta-SDT model formulation

Consider again the response *A* ∈ {1, 2, 3, 4} on our ordinal scale. Define a conditional threshold model by

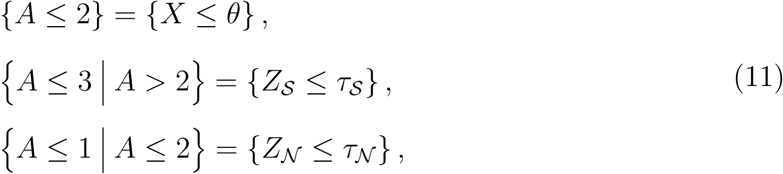

for three independent, latent variables *X*, *Z*_𝒩_ and *Z*_𝒮_. The associated distributional model is

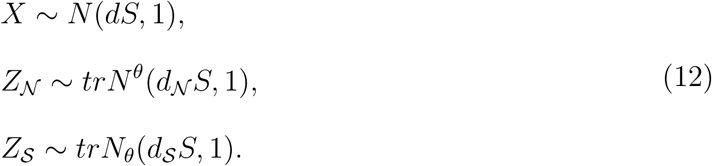

Note that the right-hand side of Figure 4 depicts the density of *Z*_𝒮_ by the fully drawn lines (black / blue for *S* = 0 and gray / red for *S* = 1). We will refer to the model specified by (11) and (12) as the meta-SDT model.

We refer to the parameters *d*_𝒩_ and *d*_𝒮_ as the response-specific meta-*d*’s and take the meta-SDT model as their formal definition. This model formulation leads to estimates of meta-*d* coincident with those obtained from the method in for example Barrett et al., 2013 where *τ*_𝒮_ and the signal-specific meta-*d* are defined as the *t*_0_ and *d*_0_, respectively, satisfying the equations

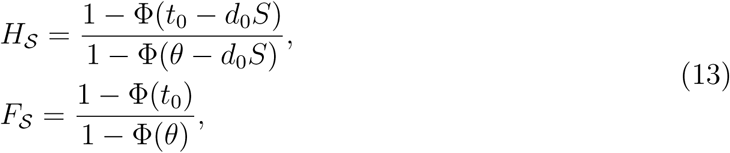

where *H*_𝒮_ is the signal-specific hit rate (the probability of reporting confidence given a correct “signal” assertion) and *F*_𝒮_ is the signal-specific false-alarm rate (probability of confidence given an incorrect “signal” response). The equations (13) are recovered in the meta-SDT model by considering ℙ (*Z*_𝒮_ > *τ*_𝒮_) and varying *S* = 1, 0 in the distribution function for the truncated normal distribution. The noise-specific meta-*d* is treated analogously.

In Barrett et al., 2013 and Maniscalco and Lau, 2014 *θ* in the type 2 task, called *θ̃*, is allowed to vary from that in the type 1 task. I.e. our approach corresponds to *θ̃* = *θ*. We return to this in the discussion.

### Meta-SDT likelihood

We begin by writing the likelihood of a multinomial distribution in a convenient form. Let *A* ∈ {1, 2, 3, 4} be a multinomial observation and suppose that the scale is ordered. The log-likelihood for the observation when parametrising the cumulative probabilities by some *ψ* is then, up to a constant, given by

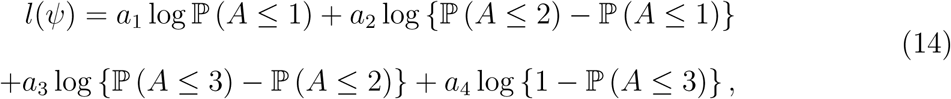

where *a_k_* = 𝟙_{*A*=*k*}_. Inserting the probabilities from (6) or (9) would respectively yield the PO or PPO model.

By conditioning, we can write the same likelihood in another manner,

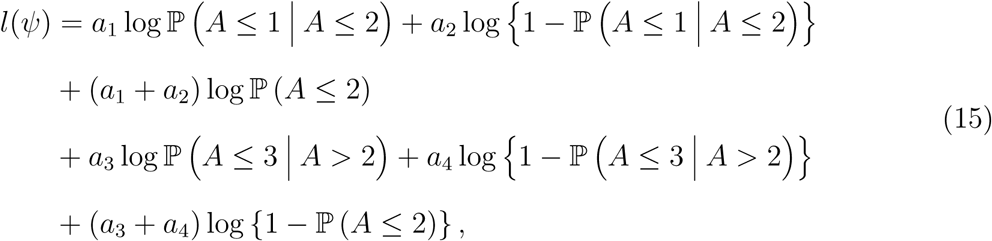

and this form immediately admits the meta-SDT model in (11) and (12). In other words, the meta-SDT is a model for the same ordinal response as SDT and we note that this in fact reduces to the standard SDT when *d*_𝒩_ = *d* = *d*_𝒮_. To perform estimation in the meta-SDT model, we would maximise (15) as a function of the parameters.

Inserting the probabilities from (11)–(12) and differentiating (15) with respect to *τ*_𝒮_ and *d*_𝒮_ we see that the parameter values maximising the likelihood must satisfy the equations in (13). In other words, in the simple meta-SDT model with a dichotomous type 2 response, the maximum likelihood estimates for *d*_𝒩_ and *d*_𝒮_ coincide with the estimators proposed in Barrett et al., 2013.

Similarly, one may verify that the estimators of *θ* and *d* will be given by (1). In other words, the type 1 parameters in the simple case meta-SDT model are estimated in the same manner as in the standard type 1 task SDT model.

### The general model formulation

We now formulate the meta-SDT model for a general response scale in a regression framework. While notationally more cumbersome the idea is the same as that considered in (11) and (12).

We consider an experiment in which a subject must perform a mutually exclusive two options choice as the type 1 task and in the type 2 task rate some aspect of the performance on an ordinal scale with 1, …, *L* levels. *A* is still a concatenation of the type 1 and type 2 tasks, so that *A* ∈ {1, …, *K*} for *K* = 2*L*, cf. Figure 5.

**Figure 5.**
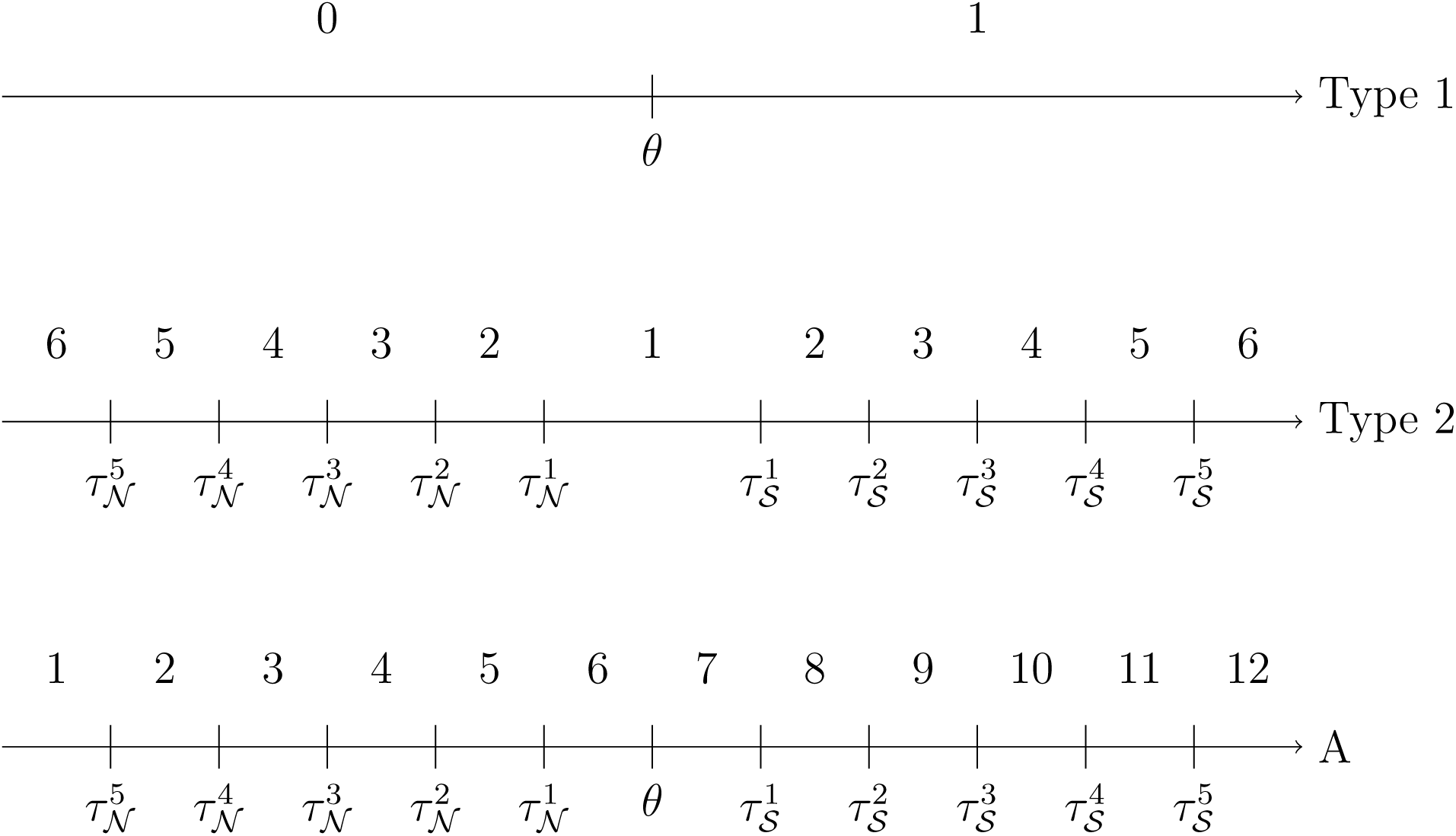
Response scales in the meta-SDT model for *L* = 6.

Define the threshold model by

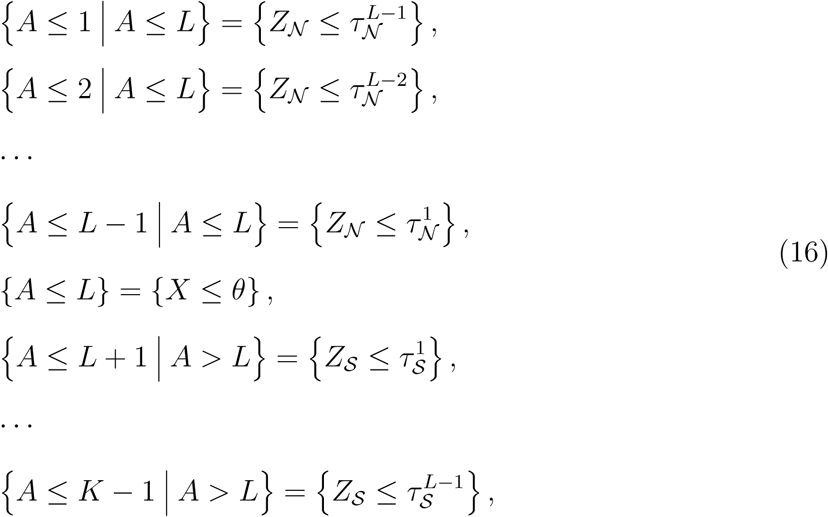

for three independent, latent variables *X*, *Z*_𝒩_ and *Z*_𝒮_. We apply the same distributional model as above but allow a more flexible mean structure,

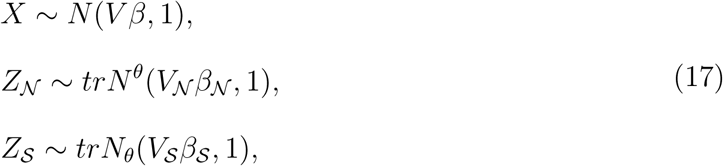

for covariate matrices *V*, *V*_𝒩_ and *V*_𝒮_ and parameter vectors *β*, *β*_𝒩_ and *β*_𝒮_.

The parameters of the model are the thresholds 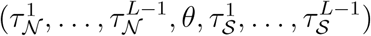 and the mean parameters (*β*, *β*_𝒩_, *β*_𝒮_), which together we will denote as *ψ*. The log-likelihood function is

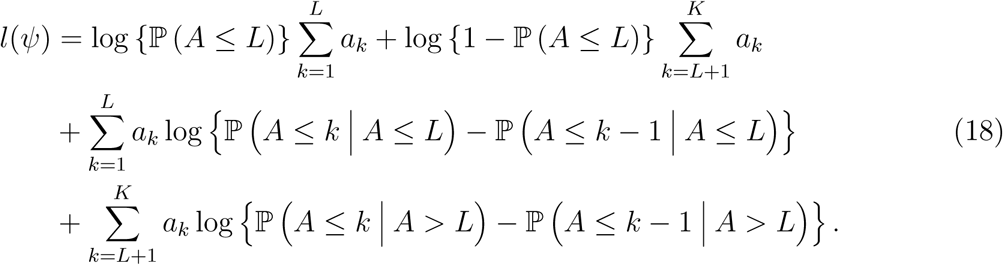

We postpone some notes on numerical aspects to the section on the `metaSDTreg` package below.

### Additional points concerning the meta-SDT model

Below we briefly present a few select points on the meta-SDT model in a discussive form.

#### Meta-SDT and the PPO model

Let us clarify the relationship between the meta-SDT model and the PPO model. For simplicity we will assume that the confidence scale only has two levels, i.e. *L* = 2, and with only one signal intensity, *S* ∈ {0,1}. We compare to the PPO model given by

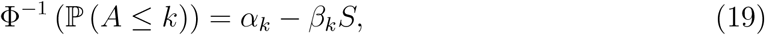

for *k* = 1, 2, 3. Note that this is in fact equivalent to three separate probit regressions on the cumulative probabilities and thus the estimates are of the form in Appendix A.

By differentiating the likelihood in (15) and studying the resulting estimating equations one may show that the intercept estimates will coincide between the two models. Therefore we will write *α*_1_ = *τ*_𝒩_, *α*_2_ = *θ* and *α*_3_ = *τ*_𝒮_. Similarly, examining the estimating equation for *d* in meta-SDT we note that this coincides with that for *β*_2_ in the PPO model and thus the two models will give the same estimate *d* = 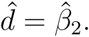

The difference between the two models lies in the manner in which confidence ratings occur. In the PPO model

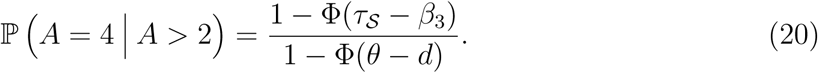

In the meta-SDT model,

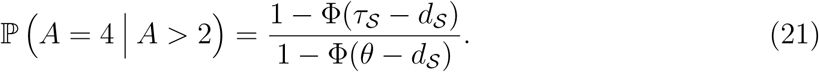

Comparing (20) and (21) we see that the difference lies in the conditional event. In the PPO model, we condition on the subject’s “signal” decision, which is performed with the sensitivity *d*. In the meta-SDT model, we condition on the hypothetical or unobserved event that the subject responded “signal” using the signal-specific meta-*d* as sensitivity. This situation arises if the subject’s type 1 and type 2 assertions were both based on a standard SDT model with sensitivity *d*_𝒮_, however the response from the type 1 task in this model is unobserved.

A consequence of this is that in the PPO model the marginal, cumulative probabilities depend only on one sensitivity parameter, while in meta-SDT it depends on two. Conversely, if one considers the conditional probabilities given the type 1 response, the meta-SDT describes this probability using one sensitivity (cf. (21)), while the PPO uses two (cf. (20)). From this, we may characterise the PPO model as focusing on the marginal probabilities where the meta-SDT focuses on the conditional. This may be seen as a consequence of the original definition of meta-*d*’ from the response-conditional ROC curves (Maniscalco & Lau, 2012).

#### Model validation

The meta-SDT model relies on what might be termed a proportional odds assumption when *L* > 2 in that the response-specific type 2 responses each depend on a single latent variable. More precisely, the latent variable does not depend on *k*.

As stated above, such assumptions are in the PO model often assessed using proportional odds plots, but the nonlinear, noninvertible nature of the response-specific probabilities renders this approach impractical for the meta-SDT model. Note however, that a proportional odds plot may still be used to validate the standard SDT model as this is equivalent to the PO model. A common technique in Signal Detection Theory is to compare the observed response-specific ROC curves to that predicted by the model. This approach, however, suffers from the disadvantage that it is difficult to interpret deviations from the predicted ROC curve. Here we will discuss a model-based approach to checking the assumptions which can be complemented by a formal test. We focus on the signal-specific response case, but the noise-specific case is symmetric.

In the meta-SDT with signal variable *S* ∈ {0,1}, the confidence in a “signal” assertion on the scale {1, …, *L*} is determined by the latent variable

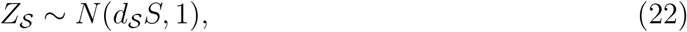

which is independent of the confidence scale level *k*. This is a special case of (17).

We may expand this model to obtain probabilities

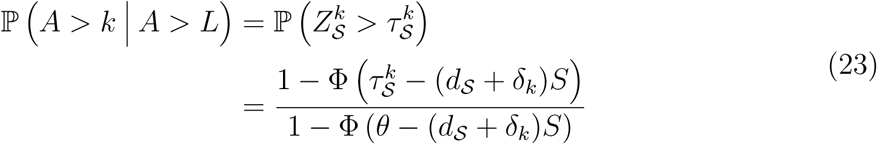

for *k* = *L* + 1, …, *K* – 1 where for identifiability we set *δ*_*L*+1_ to zero. Thus, the signal-specific meta-*d* applied by the subject to decide in a confidence of more than *k* is *d*_𝒮_ + *δ_k_*. This is an explicit model of any deviation from proportionality, the deviation being measured by the *δ_k_*’s. We will refer to this model as “saturated”.

A test that may be relevant is a general goodness of fit test to compare the predicted, marginal probabilities to those observed under the assumption that *A* ∈ {1, …, *K*} has an unstructured multinomial distribution. We discuss this test in the setting of a binary signal, *S* ∈ {0,1}, but it is easily extended to a situation where the observation of *A* over factor variables in *V*, *V*_𝒩_ and *V*_𝒮_ can be summarised in a contingency table. Suppose that we have predicted marginal probabilities for {*A* = *k*} for *S* ∈ {0,1}, say 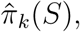 a total of 2*K* probabilities parametrised by *K* – 1 intercepts and three sensitivities, a total of *K* + 2 parameters. If the observed probabilities, say *p_k_*(*S*), come from an unstructured multinomial distribution, they must be parametrised using 2*K* – 1 parameters. The test is performed by evaluating the likelihood ratio testor

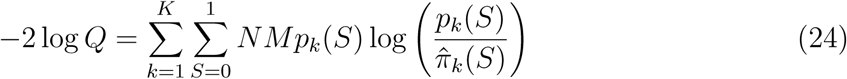

against a *χ*^2^ distribution with degrees of freedom *DF* = 2*K* – 1 – (*K* + 2) = *K* – 3. Usually this test is recommended only when the expected number of counts per cell is larger than five.

Note that the goodness of fit test in (24) will only be suggestive as the ordered model is not formally nested within the multinomial. However, this can be ammended, and indeed one would obtain a more informative test, if one were to use the expected probabilities arising from the saturated, ordered model described above. This can be achieved by fitting the model in (23) as an expansion of the meta-SDT model. This can also be implemented by applying the meta-STD model for each *l* ∈ {1, …, *L* – 1} as if one were dealing with a dichotomous type 2 rating cut at *l*.

Starting from the saturated model in (23) we may formally test the hypothesis of proportionality by testing

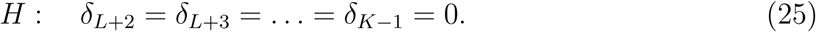

The test can be performed, for example, by evaluating twice the difference in log-likelihood between the expanded model and the unexpanded in a *χ*^2^(*DF*)-distribution with degrees of freedom *DF* = *L* – 2. Alternatively, calculating the –2 log *Q* statistic we may compare the observed probabilities to those expected under the saturated model. We can then further compare to that expected under the meta-SDT model by evaluating the deviance in a *χ*^2^(*L* – 2)-distribution. In other words, we do not need the expanded model but only the predicted probabilities.

#### A joint measure of meta-*d*

When the response-specific meta-sensitivities *d*_𝒩_ and *d*_𝒮_ are comparable, it may be desirable to aggregate these into a joint measure of metacognitive sensitivity which we will denote *d̃*. One may obtain such a measure by applying the meta-SDT model with the restriction *d*_𝒩_ = *d*_𝒮_ = *d̃*. Alternatively, *d̃* can be obtained as a weighted average of the response-specific meta-*d*’s.

In Barrett et al., 2013 it is suggested in the simple SDT to concatenate the measures as

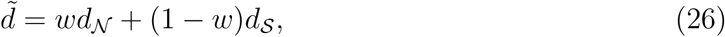

where the weight is taken to be the proportion of “noise” responses in the type 1 task.

A general approach to the same problem is to construct a weighted estimator of the form in (26) that has minimal variance (i.e. standard error) among the class of such estimators. Using that the variance of the joint measure is

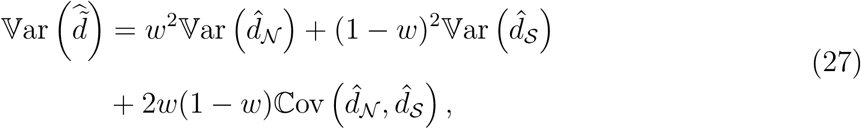

one may show that the optimal weight is

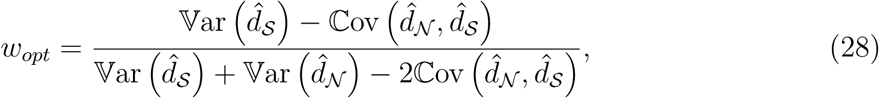

i.e. the estimators are weighted inverse of their relative contribution to the standard error. Note also that we may calculate the standard error of the associated optimal estimator of *d̃* by inserting *w_opt_* in the variance expression in (27).

This technique may be applied as a general and principled approach to obtaining a joint measure of meta-sensitivity, which can also be applied in more complex models than that considered in Barrett et al., 2013. In practice, one could insert the numerical version of the variances and covariance as obtained from the observed information, which we return to in our discussion of the `metaSDTreg` package below.

#### Approximate closed-form estimates

Here we will argue that when the latent variable distributions in the meta-SDT model are taken to be logistic rather than Gaussian, closed-form estimates for *d*_𝒩_ and *d*_𝒮_ can be obtained in the simple, dichotomous type 2 rating and binary signal, case. This can be combined with an approximation between the Gaussian and logistic cumulative distribution function to derive closed-form approximate estimates of the response-specific meta-sensitivities in the Gaussian meta-SDT model. A précis of the approximate relation between the logit and probit can be found in for example Demidenko, 2013, Section 7.1.1. Here we will apply a simple scaling approximation Φ(*x*) ≈ Λ(*cx*), where Λ is the logistic cumulative distribution function and *c* is a scaling constant (which is usually taken to be *c* = 1.7).

Applying the approximation to the signal-specific type 2 hit rate,

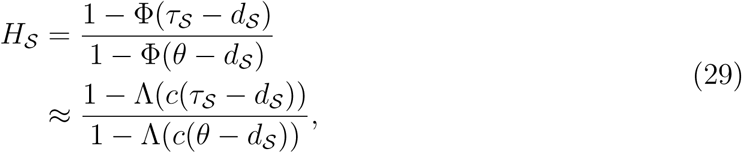

we note that (29) is equivalent to

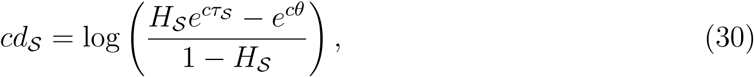

which can be used to estimate *d*_𝒮_. Specifically, suppose that *h*_𝒮_ is the observed frequency of confident answers given a correct “signal” assertion, and that we have estimators 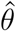 for *θ* and 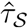 for *τ*_𝒮_. We would then estimate *d*_𝒮_ by

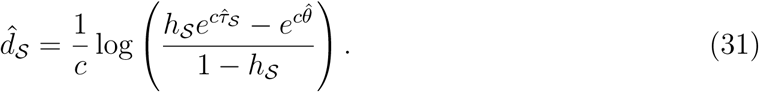

Similarly, one may derive the approximate estimator for the noise-specific case,

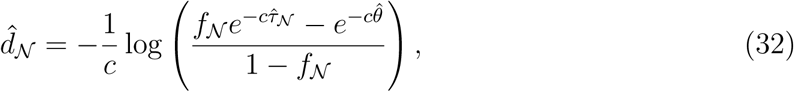

where *f*_𝒩_ is the observed, type 2 noise-specific false alarm rate.

It is of theoretical interest to study the logistic version of the estimates further. If we start from a logistic SDT model, *X* ∼ Λ, we have that

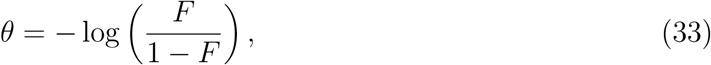

and

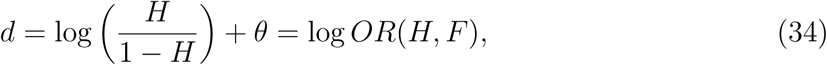

i.e. *d* is the log odds ratio comparing the odds of a hit to that of a false alarm. Defining then the, say signal-specific, type 2 model as before through the truncated logistic distribution, yields

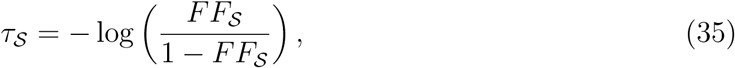

where *FF*_𝒮_ is simply the probability of reporting confidence in a “signal” assertion when the signal is absent. For the signal-specific sensitivity we obtain the same expression as in (31) with *c* =1. This can be written,

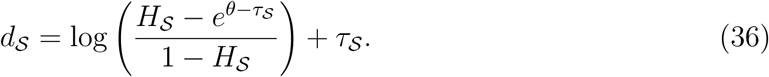

We recognise the same structure as in (34), but we are not aware of an obvious interpretation of (36). However, if *τ*_𝒮_ is much larger than *θ* so that exp(*θ* – *τ*_𝒮_) ≈ 0,

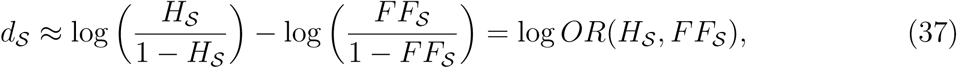

i.e. in this case, *d*_𝒮_ is approximately a log odds ratio.

## A regression model

In the present section we will consider an experiment such as that described in the introduction. The purpose is to discuss how one might approach the analysis using a regression model and additionally to, through simulation in simple cases, make some points concerning the importance of accounting for the different sources of variation in an efficient manner.

We will let *A_ij_* be the response of the *i*’th (1, …, *N*) individual in the *j*’th (1, …, *M*) replication (trial) of the experimental setting. The subject must for example claim presence or absence of a visual structure in an image to which the subject is exposed for *t_ij_* milliseconds. Let *S_ij_* ∈ {0,1} denote the presence of a stimulus in the given trial. Afterwards, the subject must rate the awareness of the type 1 decision using an *L*-point awareness scale. The combination of the type 1 assertion and the awareness rating is concatenated in the ordinal variable *A_ij_* ∈ {1, …, *K*} as previously described (see Figure 5), with *K* = 2*L*.

We will suppose that individuals prior to the experiment are randomised into one of two groups (𝓐 or 𝓑). The main purpose of the study is to estimate the overall group effect on the cognitive and meta-cognitive sensitivities of the subjects.

A simple model taking into account the above described aspects would be the general meta-SDT model in (16) and (17) taking

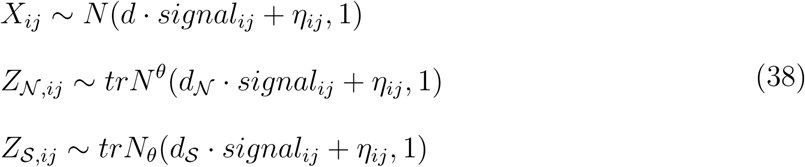

where

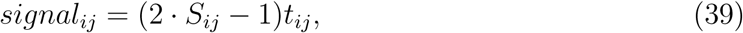

is the directed signal intensity. This ensures and assumes that the intensity of a stimulus affects equally a subjects propensity to report “noise” and “signal”.

The linear predictor in (38) is

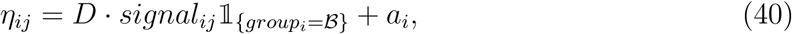

so that *D* is the effect of group 𝓑 while *a_i_* is the subject-specific response bias. These effects are assumed to enter proportionally, i.e. they are independent of *k*. Naturally, one could also have included a subject effect in the sensitivities, but we have omitted this here for simplicity.

We make a few remarks on the parametrisation. Note that the latent scale is fixed by letting zero be the mean of the underlying normal distribution when the signal intensity is zero (*t_ij_* = 0) and we are considering an unbiased subject (*a_i_* = 0). For such a trial, the subject’s probability of a “signal” assertion is 1 – Φ(*θ*), and specifically if the population is unbiased (*θ* = 0), the probability is 1/2. This corresponds to the intuition that, in the absense of bias, a subject will answer at random when presented with no stimulus. Also note that the cognitive parameters of a subject in group *𝓐* are (*d*_𝒩_, *d*, *d*_𝒮_), while the interaction between the signal and group variable means that a subject in group 𝓑 has cognitive parameters (*d*_𝒩_ + *D*, *d* + *D*, *d*_𝒮_ + *D*).

## Simulations

### Simulation study 1: Subject heterogeneity

We simulated the model in (38) with no group effect and with a dichotomous type 2 rating (*L* = 2) and a binary signal intensity (*S_ij_* ∈ {0,1}). In this case the sensitivity parameters admit estimation by using moment estimators such as those in (1) and (13) (the method in e.g. Barrett et al., 2013). The subject effects were drawn from a random distribution,

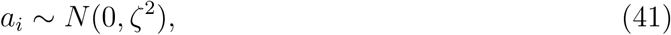

where *ζ* measures the between subject variation. In SDT this phenomenon is also referred to as “criterion jitter”, where here the jitter is on the subject level. Note that a large *ζ* corresponds to a large amount of jittering. Apart from the method of moment (MoM) estimators we additionally applied a meta-SDT regression model to account for the subject response bias. The resulting maximum likelihood estimates (MLE) were obtained using the *metaSDTreg* package. Note that although the data where simulated from a random effect model (41), the analyses applied a fixed effect model. A naïve estimate of *ζ* can in this case be obtained as the empirical standard deviation of the fixed effect estimates *â*_1_, …, *â_N_*.

We simulated the model with *N* = 25 individuals and varying the number of tasks *M* = 20, 50,100, 200. The simulations were performed for three degress of subject heterogeneity *ζ* = 0, 0.5, 0.75. Each scenario was simulated 1000 times. The resulting sensitivity estimates 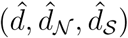 are shown in Figure 6 (**A**)-(**C**), one panel for each parameter.

**Figure 6.**
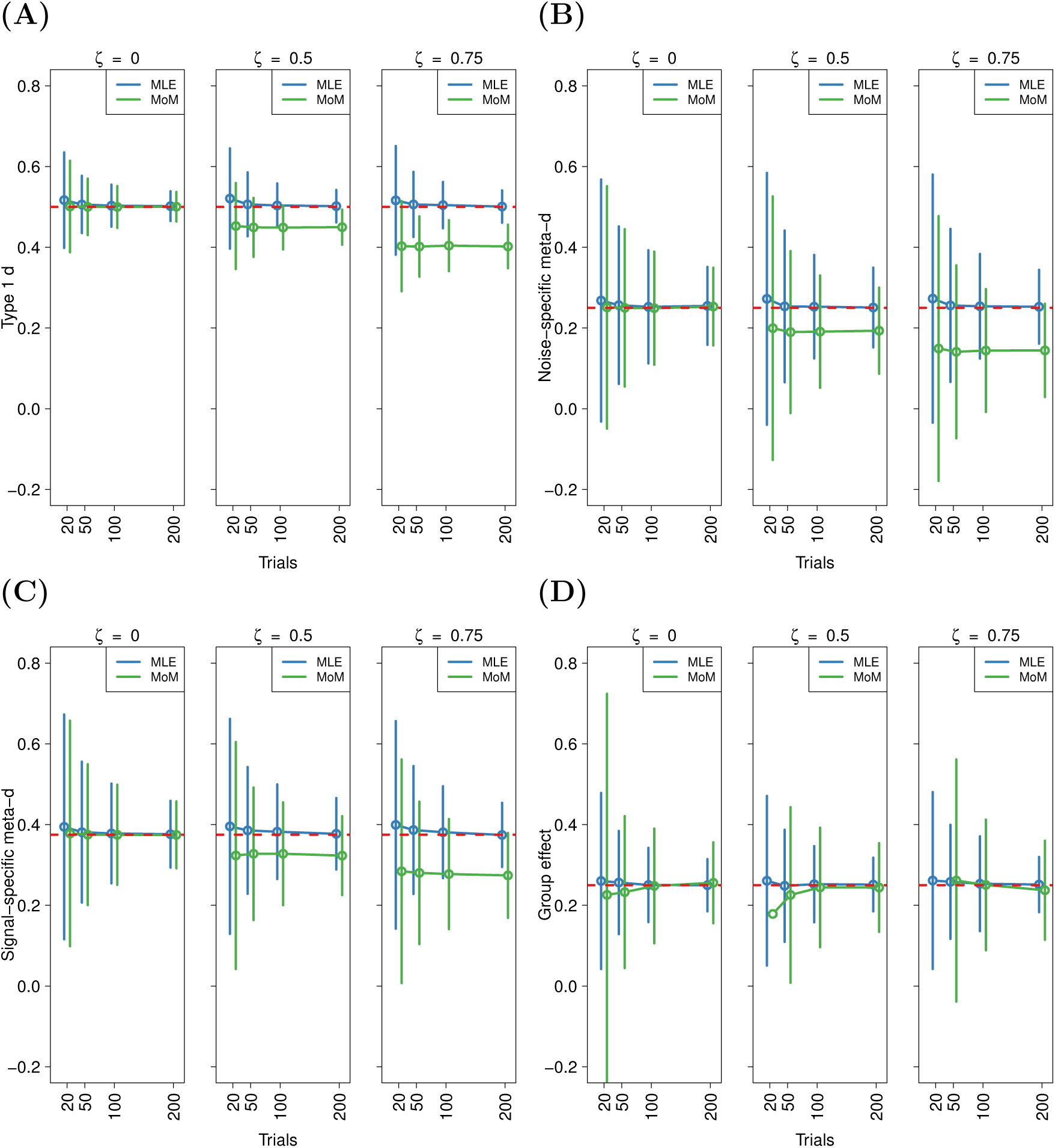
**(A)-(C)** Simulations to illustrate the effect of heterogeneity on estimates of sensitivities. For each of the three parameters, the left column is simulations from a scenario with no subject heterogeneity (*ζ* = 0) while the middle column has an intermediate between subject variation (*ζ* = 0.5) and the right column has a high degree of heterogeneity (*ζ* = 0.75). **(A)** depicts estimates of *d*, **(B)** estimates of *d*_𝒩_ and **(C)** shows estimates of *d*_𝒮_. **(D)** Estimates of group effect *D* for three levels of heterogeneity. For all graphs, the points represent simulation means and error bars are twice the empirical standard deviation of the simulated estimates (i.e. the observed standard error), while the horisontal dashed line is the true value.

We observe that in the independent subject case (*ζ* = 0), the performance of MoM and MLE is interchangeable as would be expected. However, as the between subject variation grows larger, the bias of the MoM estimates becomes more pronounced. This fact is a general lesson from non-linear and generalised linear models: While in the linear case, neglecting to account for a random effect will usually only effect efficiency of the estimator, the disregard of a random effect in a non-linear model leads to biased, inconsistent and inefficient estimation (e.g. Demidenko, 2013).

We additionally note in connection with Figure 6 that the MLE for these specific choices of parameters converge satisfactorily to the true value for *N* = 25 fixed as the number of replications per subject is increased. This will not generally hold and one would need to also increase the number of subjects (Demidenko, 2013) to ensure convergence to the true value. However, to study the asymptotics of the estimates as *N* grows to infinity one would need a random effect model as in our present, fixed effect, approach the number of parameters will grow with *N* leading to inconsistent estimates.

### Simulation study 2: Efficient group comparisons

One way to amend the performance within the method of moments is instead to apply the MoM estimators on a subject level, as is often done. This is however inefficient and requires a large number of replications per subject, *M*, as the subject must have responses in every cell of the (*A*, *signal*) table. Often this leads to exclusion of some individuals resulting in another form of bias in the estimation process. We explore this subject level analysis in the simulations below.

We now simulate the model in (38) for the same three degrees of subject heterogeneity as in Study 1 but with a group effect *D* = 0.25 and with a dichotomous type 2 rating and a binary signal intensity. We assume that apart from estimating the group parameter, one wishes to adjust for any subject effect. Thus, for all simulations, *N* = 25 individuals and we vary the number of trials *M* = 20, 50,100,200 and heterogeneity *ζ* = 0, 0.5, 0.75. Again, each scenario was simulated 1000 times We compare the regression model with a subject-specific intercept fitted by maximum likelihood against the MoM estimates obtained by applying the analysis on a subject level and then regressing the obtained sensitivity parameters against *group* to obtain an estimate of *D*.

The results of Simulation study 2 is plotted in Figure 6, panel **(D)**. As expected, the error bars are more narrow for the MLE estimates as it is more efficient to obtain the group estimate from an overall model. More importantly, we see that for low number of replicates the MoM estimates are biased. When subject heterogenity was present, *M* = 20 trials proved insufficient to obtain sufficient sensitivity estimates. Thus, for *ζ* = 0.5 only one analysis resulted in an estimate of *D* (and as such, the error bar could not be calculated), while no estimates could be otained for *ζ* = 0.75.

The observed bias is due to the exclusion of subjects when a sensitivity parameter can not be determined showing that subject level analyses do not merely lead to imprecision but also to bias.

## The metaSDTreg package

The procedure in `metaSDTreg` proceeds in two steps by first calculating the PPO estimates as detailed above and then maximising (18) with PPO estimates as starting values. The negative Hessian associated with the procedure is the numerical version of *j*(*ψ*) = –*∂*^2^*l*(*ψ*)/*∂ψ∂ψ*^*T*^, that is, the observed information of the model, which can be viewed as an empirical estimate of the asymptotic, inverse covariance matrix of 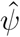. Thus, it can be used to construct approximate confidence intervals and tests. An alternative would be to use the Fisher information 𝔼 [*j*(*ψ*)] to perform inference. The Fisher information could also be applied directly in the optimisation (so-called Fisher scoring) with the advantage that it is always non-negative definite in a properly specified model in contrast to the observed information matrix (Demidenko, 2013, section 2.11).

## Discussion

In the section below we will delimit our present approach with regards to similar methods and we discuss possible directions for future developments.

### Choice of type 1 threshold in the response-specific type 2 model

In the literature on meta-*d* it is suggested to consider another type 1 threshold than *θ* for the type 2 models. We would thus have a *θ̃*_𝒩_ in the noise-conditional type 2 model and *θ̃*_𝒮_ in the signal-conditional type 2 model. In our notation, this would mean that *Z*_𝒩_ should follow a *trN*^θ̃_𝒩_^-distribution and *Z*_𝒮_ a *trN*_*θ̃*_𝒮__-distribution. We proceed with the discussion in the signal-specific case.

Naturally, as *θ̃*_𝒮_ is the type 1 threshold in the unobserved part of the signal-specific SDT (see the section “Meta-SDT and the PPO model” above) it can not be estimated, and one will generally need to fix it as some function of the quantities *θ*, *d* and *d*_𝒮_. Barrett et al., 2013 proposes to use the relative type 1 threshold *θ̃*_𝒮_ = *θd*_𝒮_/*d*. Maniscalco and Lau, 2014 suggest the same measure as well as one based on the likelihood ratio.

Interestingly, if we would allow the function to also depend on *S*, we could take *θ̃*_𝒮_ = *θ* + (*d*_𝒮_ – *d*)*S* and we would recover the PPO model (cf. equation (20) and (21)). We can extend this argument and apply the approximation log *z* ∼ *z* – 1 formally to show that if we take *θ̃*_𝒮_ to be the relative type 1 threshold, then

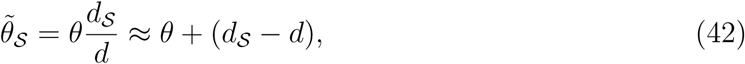

so that for *S* =1 the meta-SDT signal-specific hit rate is approximately the same as that in the PPO.

### Random and scale effects

In our expansion of the meta-SDT in (17) we focused on location effects, i.e. effects that shift the mean of the latent variables. We could also have expanded the variance structure on the latent variables and allowed this to depend on covariates leading to scale effects. A common scale effect in signal detection theory is found in the Unequal Variance SDT model, which in statistics is known as a heteroscedastic probit model (e.g. DeCarlo, 2010), where the variance of the latent variable is allowed to depend on the presence or absence of a signal.

Additionally, it would be desireable to include random effects, as we have already explored in the simulations. However, estimation procedures for random effects are not implemented in the `metaSDTreg` package at the time of writing, as the likelihood function in this case contains an intractable integral and approximative procedures such as Gaussian quadrature or Laplace approximation are needed (Demidenko, 2013, section 7.1-7.2).

### Closed-form estimates

Another aspect that might be worth investigating is the applicability of the closed form estimates in (31) and (32). Specifically, even though the error between the cumulative distribution functions of the probit and logit is small (Demidenko, 2013), it would be interesting to investigate the error on the meta-*d*’ estimates. Moreover, it may be possible to improve the approximation for example by focusing on an approximation of the truncated distribution function rather than the distribution function itself.

Apart from the theoretical value of such an investigation, precise, closed-form estimates may prove a robust manner of obtaining starting values for more advanced procedures.

### Indices of metacognitive abilities

It has been suggested that when quantifying metacognitive ability using the sensitivities *d*, *d*_𝒩_ and *d*_𝒮_, one should evaluate the metacognitive sensitivities in comparison to the type 1 sensitivity (Maniscalco & Lau, 2012, 2014). Both the absolute difference *d*_𝒮_ – *d* and the relative difference *d*_𝒮_/*d* have been proposed.

If we think of meta-*d* as a log-odds ratio, cf. (37) and also (34), this would suggest that one should consider the absolute difference, as this could be interpretated as a log-odds ratio ratio, while the relative difference would have a less straightforward interpretation.

Considering the simulation results in Figure 6 **(A)** - **(C)** one might suspect that while the sensitivities are biased separately, the index might retain smaller bias. We found this to hold for the absolute difference for all choices of *ζ* but less so for the ratio. Of course, we caution against interpreting this as a argument for the absolute difference over the ratio or that bias in general can be avoided by considering absolute differences, as the results are based on a single, one-case, simulation study.

### Serial correlation

An aspect that may need to be investigated in future developments is that of serial correlation, although this is generally challenging in non-linear models (e.g. Breinegaard, Rabe-Hesketh, & Skrondal, 2015).

In our approach, we assume that the replications on a single subject may be assumed to be independent given the covariates (in particular the subject’s threshold and sensitivity). However, in many cognitive experiments it is easy to imagine that observations taken close together in time could be more correlated than observations far apart, in which case we say that the correlation is serial. Indeed, one might imagine negative serial correlation to occur if a subject were adverse to give a response having already given the same response multiple times in a row. Conversely in a visual study, the presence of a structure in an image might leave a “residue” that would prompt the subject to claim the structure as present in the following trial, thus leading to positive serial correlation.

### Bayesian approaches

Recently there has been some effort in defining hierarchial SDT models and hierarchial models for meta-*d* in a Bayesian framework. For example, Fleming, 2017 discusses the implementation of a model for meta-*d* in a Matlab toolbox, which is a front-end to the MCMC sampler JAGS. We are also aware of similar approaches using the MCMC program STAN through the R-package `brms` (Bürkner, 2017), although we do not know of an attempt to study meta-*d* using this package.

The main difference between Fleming, 2017 and the approach of the present paper is that Fleming, 2017 starts from the multinomial type 2 likelihood as found in Maniscalco and Lau, 2014. Additionally, we are not aware that the approach offers regression capabilities. This is in contrast to the meta-SDT model which is a model for all the responses arising from the cognitive experiment, formulated through latent variables which is well-suited for expansions to include covariate information. Another difference is the manner in which inter-subject variation is modelled, where our approach directly parametrises this variation as effects and where the Bayesian approach in Fleming, 2017 uses hyperpriors to incorporate the extra variation.

There is no reason that the model we proposed in (16)–(17) could not be implemented in a Bayesian manner for example in `R` through the `brms`-package. This would entail some considerations, which may be said to apply to Bayesian data modelling in general: namely, the choice of prior distributions and methods for assessing convergence of the Markov chain. Moreover, the importance of these considerations become increasingly critical in non-linear models as these often demand more informative priors to converge (as also discussed in Bürkner, 2017). Thus, one would require prior knowledge of some or even all the parameters in the regression model, which may be difficult to obtain as the interpretation of each parameter would change with the inclusion of other covariates.

As a tentative way of comparing the methods, we applied the out-of-the-box response-specific, group level analysis of Fleming, 2017 to some simulations from Simulation study 1 above. The details and results are presented in Appendix C. These results showing better performances of the MLE procedure over HMeta-d should naturally be interpreted very cautiously as they represent simulations under a very specific scenario. A more realistic delimitation of the performance of the various methods will have to be investigated in a large-scale simulation study, which is beyond the scope of this article.

## Summary

We have presented a statistical model for metacognitive sensitivity, the meta-SDT, and have shown how it is well suited for expansion. Specifically, we can derive regression methods for metacognitive experiments. We have also shown that the standard signal detection theory model is equivalent to the proportional odds model, which is a widely used regression technique available in standard statistical software. We have made some theoretical points concerning the meta-SDT model and compared it to the partial proportional odds. Also, we supplied formulas for closed-form approximate estimates of meta-sensitivity and derived an optimal weight to combine response-specific measures. Finally, we described the implementation of the meta-SDT model in the `R`-package `metaSDTreg`, which is available on CRAN.

The meta-SDT model constitutes a theoretical contribution to signal detection theory and the theory on meta-*d*. It allows researchers to build flexible and efficient models for metacognitive experiments, and its implementation is useful for practical researchers of metacognition.

## Appendix A

### ML Estimation in probit regression

In the present appendix we derive the maximum likelihood estimators of a probit model with a binary explanatory variable. The results may be standard especially to readers familiar with the theory of exponential families, but we take a more hands on approach and study the likelihood directly.

#### Model and likelihood

Suppose that we have *N* Bernoulli observations

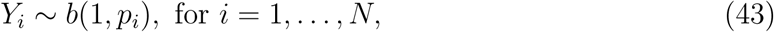

and say that we wish to model the probability parameter in a GLM with probit link, that is

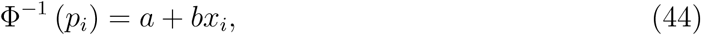

where *x_i_* ∈ {0,1}. Write *η_i_* = *a* + *bx_i_*. We wish to estimate *a* and *b* based on data such as that given in Table A1.

**Table A1.** Data for the Bernoulli observations Y and the binary, explanatory variable x.

|  |  | y |  |
| --- | --- | --- | --- |
|  |  | 0 | 1 |
| x | 0 | $n_{00}$ | $n_{01}$ |
| | 1 | $n_{10}$ | $n_{11}$ |

The log likelihood of the model is given by

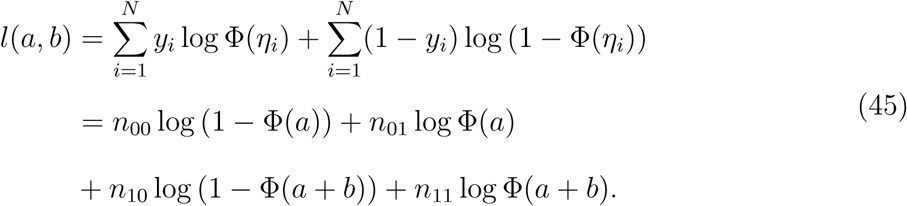

#### Likelihood equations

Differentiating the expression in (45) we obtain

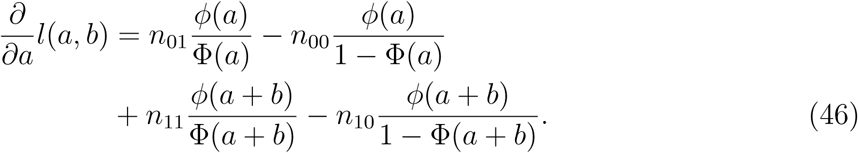

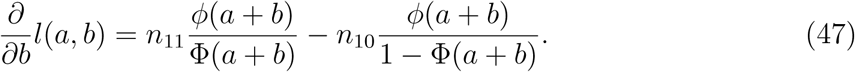

It is straightforward to verify that the likelihood equation 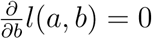 is satisfied for

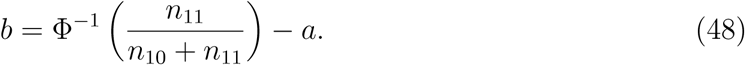

Taking the right hand side of (48) as *b̂* and plugging this into (46) we see that the corresponding equation for *a* is

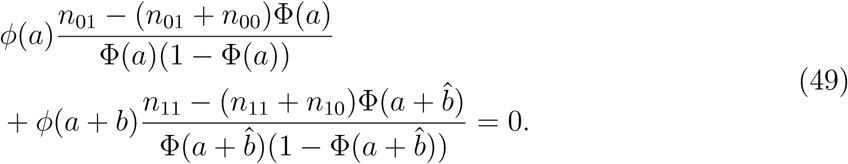

Noting that

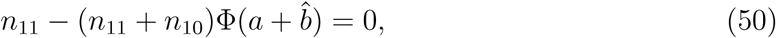

the likelihood equation for *a* reduces to

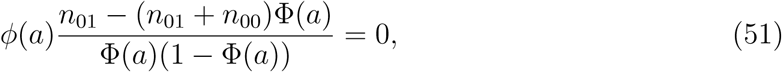

which is seen to be solved for

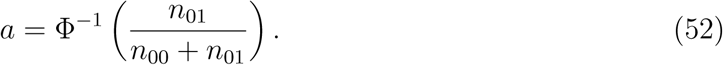

In conclusion,

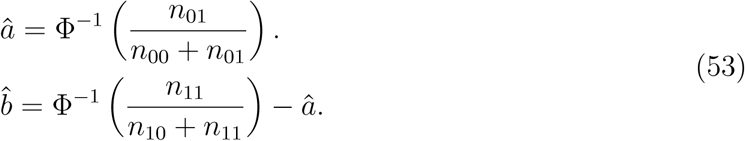

## Appendix B

### The truncated normal distribution

Recall that a stochastic variable *V* is said to follow a truncated normal distribution with lower threshold *a*, upper threshold *b*, mean *μ* and variance *σ*^2^ if

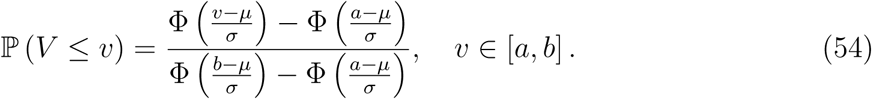

For such a variable we write 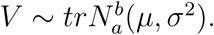 Note that if 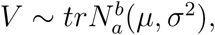 then *V* ∼ *U* | *a* ≤ *U* ≤ *b* for some variable *U* ∼ *N*(*μ*, *σ*^2^) which motivates the distribution’s name. We have that

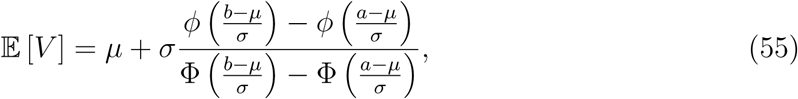

see for example Johnson, Kotz, and Balakrishnan, 1994.

Since 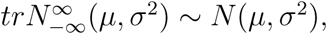 we use the notational convention of omitting a threshold when it is trivial. More precisely, we write 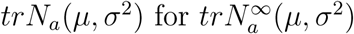 and 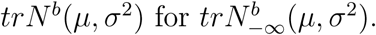

## Appendix C

### Additional simulations

We simulated 100 cases as in Simulation study 1, but now only varying the between subject variation as *ζ* = 0 and *ζ* = 0.75. For each simulation, we applied the meta SDT regression model as in Simulation study 1. Additionally, we analysed the data using the response-specific, group level analysis from the *HMeta*-*d* toolbox. All priors and MCMC initialisation parameters were unaltered from the “out of the box” settings in the toolbox. The results are shown in Figure C1.

**Figure C1.**
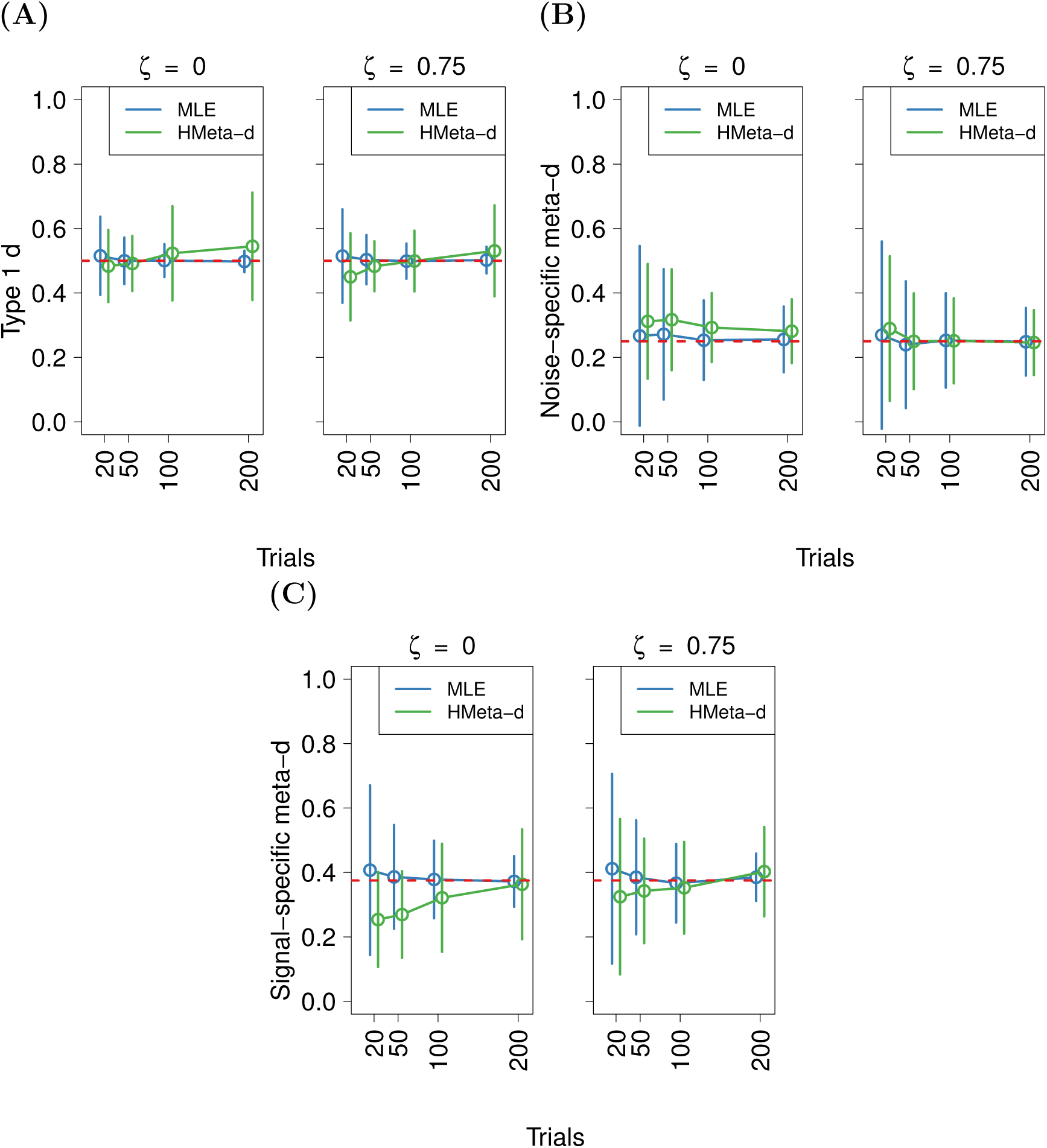
**(A)-(C)** Simulations to tentatively compare to the performance of the *HMeta*-*d* toolbox. For each of the three parameters, the left column is simulations from a scenario with no subject heterogeneity (*ζ* = 0) and the right column has a high degree of heterogeneity (*ζ* = 0.75). **(A)** estimates of *d*, **(B)** estimates of *d*_𝒩_ and **(C)** estimates of *d*_𝒮_. For all graphs, the points represent simulation means and error bars are twice the empirical standard deviation of the simulated estimates (i.e. the observed standard error), while the horisontal dashed line is the true value.

The results show that in this specific case the MLE outperforms the HMeta-d method both in terms of bias and efficiency (the width of the confidence intervals), especially at moderate sample sizes. It seems that the HMeta-d model fares better in the presence of heterogeneity compared to the case *ζ* = 0 which may be explained by the presence of hyperpriors in the model which are in a sense unused when *ζ* = 0. As already stated, this should be interpreted very cautiously as it is the result under one specific set of assumptions. In particular, as the HMeta-d model relies on a number of informative priors one would expect the performance to depend strongly on their validity in the simulated data.

